# Assessing the performance of different approaches for functional and taxonomic annotation of metagenomes

**DOI:** 10.1101/522292

**Authors:** Javier Tamames, Marta Cobo-Simón, Fernando Puente-Sánchez

## Abstract

Metagenomes can be analysed using different approaches and tools. One of the most important distinctions is the way to perform taxonomic and functional assignment, choosing between the usage of assemblies or the direct analysis of raw sequence reads instead. Many instances of each approach can be found in the literature, but to the best of our knowledge no evaluation of their different performances has been carried on, and we question if their results are comparable. We have studied this point by analysing several real and mock metagenomes using different methodologies and tools, and comparing the resulting taxonomic and functional profiles. Our results show that database completeness is the main factor determining the performance of the methods relying on direct read assignment either by homology, k-mer composition or similarity to marker genes, while methods relying on assembly and assignment of predicted genes are most influenced by sequencing depth, that in turn determines the completeness of the assembly. Although differences exist, taxonomic profiles are rather similar between raw read assignment and assembly assignment methods, while they are more divergent for methods based on k-mers and marker genes. Regarding functional annotation, analysis of raw reads retrieves more functions, but it also makes a significant number of over-predictions. Assembly methods are more advantageous as the size of the metagenome grows bigger.

## Introduction

Since its beginnings in the early 2000s, metagenomics has emerged as a very powerful way to assess the functional and taxonomic composition of microbiomes. The improvement in high-throughput sequencing technologies, computational power and bioinformatics methods have made metagenomics affordable and attainable, increasingly becoming a routine methodology for many laboratories.

A metagenomics experiment consists of a first wet-lab part, where DNA from samples is extracted and sequenced, and a second *in silico* part, where bioinformatics analysis of the sequences is carried out. There is not a golden standard for performing metagenomic experiments, especially regarding the bioinformatics used for the analysis. The usual goal of metagenomics is to provide functional and taxonomic profiles of the microbiome, that is, to know the abundances of taxa and functions.

Usually, one of the first steps in the analysis involves the assembly of the raw metagenomic reads after quality filtering. The objective is to obtain contigs, where genes can be predicted and then annotated, usually by means of comparisons against reference databases. It is reasonable to think that the taxonomic and functional identification is more precise having the full gene than just a fragment of it. Also, taxonomic classification benefits of having contiguous genes, because since they come from the same genome, unannotated genes can be ascribed to the taxon of their neighbouring genes. Therefore, the assembly can facilitate significantly the subsequent annotation steps. However, *de novo* metagenomic assembly is a complex task: the performance of the assembly is dependent on the sequencing depth and the intrinsic complexity of the microbiome [1], and it often requires large computational capacity in terms of memory usage, although recent assemblers have reduced very much this constraint. Highly diverse microbiomes, such as those of soils, are harder to assemble, likely to produce more misassembles, and will produce smaller contigs. In addition, a fraction of reads will remain unassembled, which is often considered as “losing sequences”, even if these unassembled reads are still kept and can be recovered and analysed separately. Also, many different assemblers are available, which make use of diverse algorithms and heuristics, and hence produce different results. Finally, the assessment of the quality of these assemblies is often problematic, and sometimes a high degree of chimerism can be present [2].

Because of these problems, some authors prefer to skip the assembly step and proceed to the direct functional/taxonomic annotation of the raw reads, especially when the aim is to obtain a functional or taxonomic profile of the metagenome [3–8]. This approach provides counts for the abundance of taxa and functions based on the similarity of the raw reads to corresponding genes in the database. There are two main drawbacks of working with raw reads: first, since it is based on homology searches for millions of sequences against huge reference databases, it usually needs large CPU usage, especially taking into account that for taxonomic assignment the reference database must be as complete as possible to minimize errors [9]; and second, the sequences are often too short to produce accurate assignments [10, 11].

Other methods using raw reads trust the similarity of oligonucleotide composition of sequences from the same genomes. These methods count k-mer frequency of the raw reads, and compare it to a model trained with sequences from known genomes. For instance, Kraken2 [12] or Centrifuge [13] use this k-mer counting approach. These methods can be used only for taxonomic assignment.

Also for taxonomic profiling, other methods rely on the identification of phylogenetic marker genes in raw reads to estimate the abundance of each taxa in the metagenome, for instance Metaphlan2 [14]. These methods must be considered profilers, since they do not attempt to classify as many reads as possible, but instead recognizing the identity of particular marker genes to infer community composition from these.

These different approaches (assemblies, raw reads, k-mer composition and marker gene profiling) are likely to produce different results. But we have scarce information on how different these results are, and whether they are so different as to compromise the subsequent biological interpretation of the data. This is a relevant point, since both approaches are being used indistinctly for metagenomic analyses and their results could not be comparable if the differences are large.

The objective of the present analysis is to estimate the differences between all these approaches. To this end, we classify functionally and taxonomically several real and mock metagenomes using either the raw reads or the genes derived from the assembly. The assemblies are done using the Megahit assembler [15]. For assignment of raw reads, we use Diamond a high-performance tool for homology identification that allow very fast searches [17]. For phylogenetic analysis, we also use Kraken2 as a k-mer classifier, and Metaphlan2 as a marker gene classifier.

The mock communities of known composition can help us to evaluate the goodness of the results. Even if mock communities are rather less complex than real ones, they are valuable tools for having a framework to compare the annotations done by different methods to the real expectations

We aim to illustrate how different approaches can lead to diverse results, and therefore different interpretations of the underlying biological reality. We hope that this can help in the informed choice of the most adequate method according to the particular characteristics of the dataset.

## Methods

### Overview of the procedure

We have used three different metagenomes: 1) a microbial mat metagenome from a hot spring in Huinay (Chile), corresponding to a sample taken at 48°C, and sequenced using Illumina HiSeq (82.7 M reads, 9.8 G bases, accession SRP104009) [18] 2) A marine metagenome corresponding to Malaspina sampling expedition, taken at 3 meters depth in the Pacific Ocean [19], also sequenced with Illumina HiSeq (168.9 M reads, 16 G bases). 3) A gut metagenome from the human microbiome project [20], sequenced with the Illumina Genome Analyzer II (68.1 M reads, 6.4 G bases, accession SRS052697).

A schematic description of the procedure can be seen in Figure 1. The taxonomic classification of the raw reads (from now on, RR) was obtained by direct homology searches against GenBank NR database (release 223, December 2017) using Diamond (v0.9.13.114) with a maximum e-value threshold of 1e-03 [17]. The annotations were done using a last-common ancestor (LCA) algorithm. LCA first select the hits having at least 80% of the bitscore of the best hit and overcoming the minimum identity threshold set for a particular taxonomic rank (85, 60, 55, 50, 46, 42 and 40% for species, genus, family, order, class, phylum and superkingdom ranks, respectively)[21]. This means that in order to classify a sequence at the phylum taxonomic rank, for instance, hits for that sequence must be at last 42% identical. Then it looks for the common taxon for all hits at the desired taxonomic rank (although some flexibility is allowed, for instance admitting one outlier if the number of hits is high). In case that a common taxon is not found, the read is unassigned. For the functional annotation of the raw reads (F_RR), KEGG [22] was used as the reference functional classification, and the reads were annotated using the best hit to this KEGG database.

**Figure 1:**
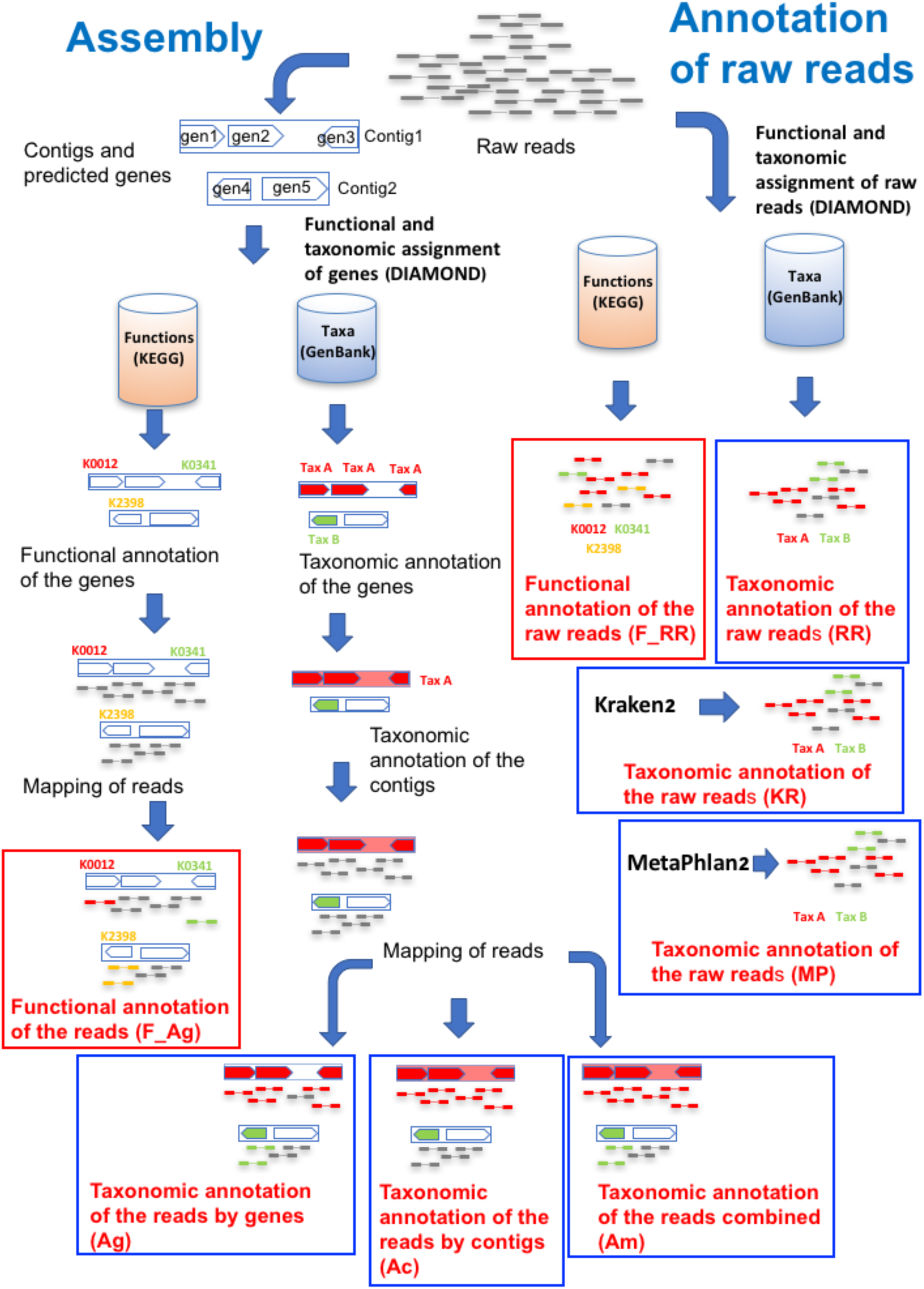
Schematic description of the procedure followed for the analysis. Boxed in blue, taxonomic annotations. In red, functional (KEGG) annotations.

The set of reads was also annotated using the assembled metagenome. We used the SqueezeMeta pipeline [23] for this task, that automatizes all steps of metagenomic analysis. The assembly was done using Megahit (v1.1.2)[15], followed by gene prediction using Prodigal (v2.6.2)[24]. The predicted genes were searched for homologies against GenBank NR and KEGG databases using Diamond, and processed with the LCA algorithm, as above. This produces taxonomic and functional assignments for the genes in the contigs.

A taxonomic classification for the whole contig can be obtained from the consensus of the annotations of its genes. The criteria for declaring a consensus taxon are: 50% of the genes in the contig must belong to the taxon, and 80% of the annotated genes (some genes may not have annotation). Otherwise, the contig is left unassigned. This approach has the advantage of allowing the annotation of many additional genes (those in the contig that were not classified directly, including orphans), but it has the drawback of dropping the original annotations for the genes if a consensus is not reached. Notice that under these criteria, short contigs comprising just one gene receive the annotation of their only gene.

For taxonomically classifying the reads, these were mapped against the contigs using Bowtie2 (v2.2.6) [25], and inherited the annotation of the corresponding gene or contig. Also, we investigated a combined approach merging these two strategies, in which reads are annotated by contigs first, and then by genes if contigs did not provide an annotation. These approaches will be referred as annotation by assembly/genes (Ag), assembly/contigs (Ac), and assembly/combined (Am). For functional classification, only mapping against genes was used (F_Ag), since there is not contig annotation for functions (each gene has a different function).

We also used two other approaches widely used in metagenomic analysis for taxonomic assignment. First, assignment by means of k-mer composition using Kraken2 (KR) [12]. Second, the clade-specific gene marker searching of Metaphlan2 (MP) [14]. These methods are not suitable for functional assignment.

For each metagenome, we compiled tables with the taxonomic or functional assignment of each of the reads by all methods. These tables were used to calculate the functional and taxonomic profiles that were used in the comparison.

For assessing the performance of the approaches, we used mock metagenomes of different sizes (2K, 5K, 1M, 2M, 5M reads) built with genomes of species significantly associated to each of the three environments considered: marine, thermal and gut. We calculated the associations between species and environments as in [26]. We selected sets of 35 environment-associated species with complete genomes available (Supplementary table 1), and calculated their abundance ratios following a hypergeometric distribution. Knowing these ratios and the total number of reads, we estimated how many reads of each species must be taken and created the mock metagenome by simulating the sequencing of the required number of paired-end reads from these genomes.

For analysing the mock metagenomes, we followed the same approaches above, but we removed the corresponding genomes in the NR database used for homology searching. We also created custom databases for Kraken2 and Metaphlan2 in which we also removed these genomes.

## Results

We analyzed three different metagenomes coming from different environments: a thermal microbial mat metagenome from a hot spring in Huinay (Chile), a marine sample from the Malaspina expedition, and a gut metagenome from the human microbiome project (thermal, marine and gut from now on). We used different methods to taxonomically assign the reads from these metagenomes (see methods for full details): 1) We ran a homology search of the reads against the GenBank NR database, followed by assignment using the last common ancestor (LCA) of the hits. We termed this approach “assignment to raw reads” (RR). 2) We used the SqueezeMeta software [23] to proceed with a standard metagenomic analysis pipeline: assembly of the genomes using Megahit [15]; prediction of genes using Prodigal [24]; Taxonomic assignment of these genes by homology search against the GenBank nr database, followed by LCA assignment as above; Taxonomic assignment of the contig to the consensus taxon of its constituent genes; mapping of the reads to the contigs using Bowtie2; and annotation of the reads according to the taxon of the gene (assembly by genes, Ag) or contig (assembly by contigs, Ac) they were mapped to. We also used a combined approach in which the read was annotated to the taxon of the contig if it is annotated, and to the taxon of the gene otherwise (assembly combined, Am). 3) We used Kraken2, a k-mer profiler that assigns reads to the most likely taxon by compositional similarity. 4) We used Metaphlan2, which attempts to find reads corresponding to clade-specific genes to assign the corresponding read to the target clade.

The results of the annotation experiment can be seen in Figure 2, for assignments at phylum rank. The results at family taxonomic rank are shown in Supplementary Fig 1, and show similar trends.

**Figure 2:**
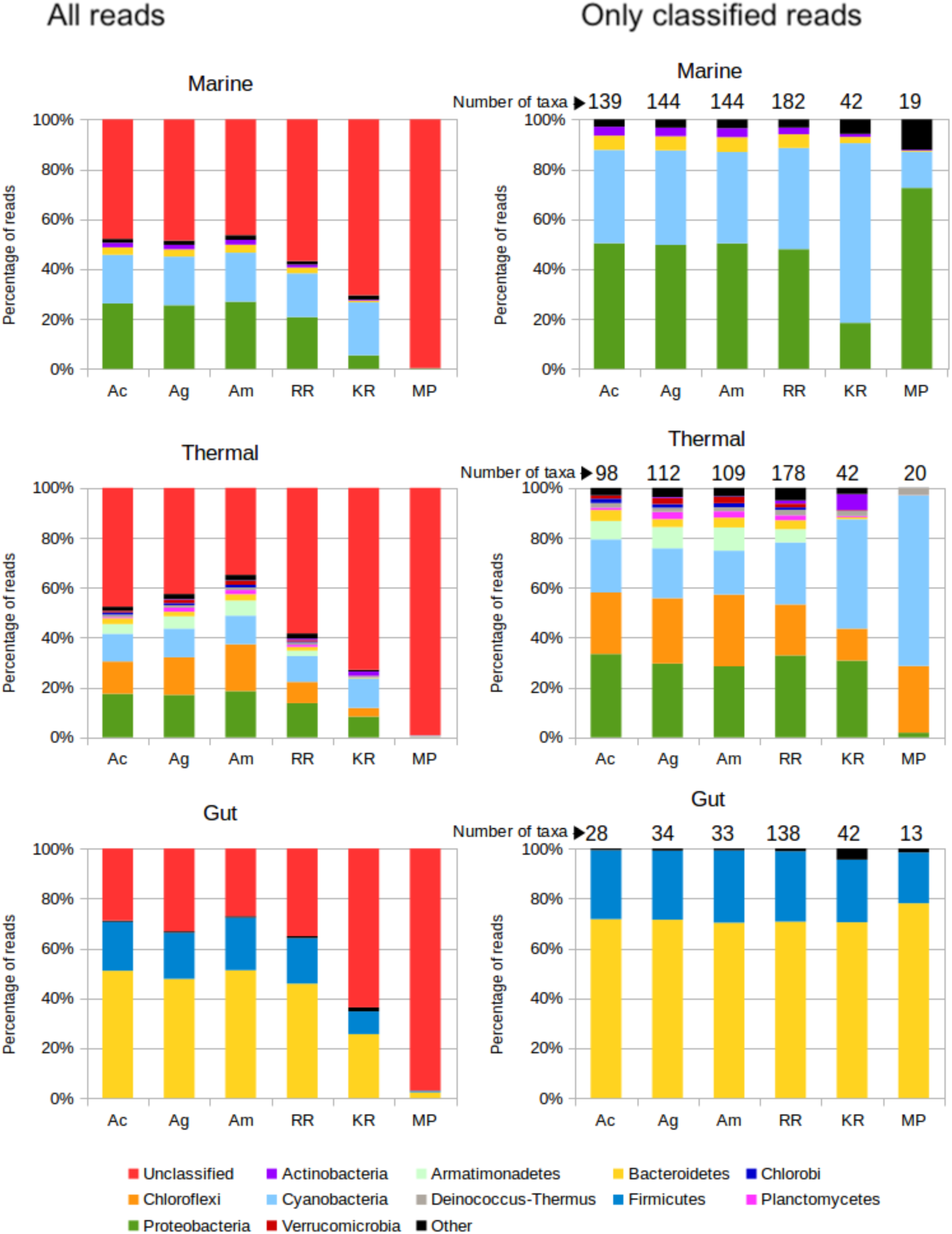
Comparison of read assignments by different methods: Ac, Assembly and mapping reads to contigs. Ag, Same but mapping reads to genes. Am, same but mapping genes first to contigs, then to genes. RR, raw reads assignment. KR: Kraken. MP: Metaphlan2. Left: All reads considered. Right: Discounting unclassified reads. These panels also show the number of obtained phyla.

Assembly methods achieve the highest number of classified reads in the three metagenomes. We anticipated that the number of classified reads by these methods would be related to the completeness of the assembly, that is, the percentage of total reads that were assembled. This will be ultimately related to the total size of the metagenome and the diversity of the community. The ratio of mapped reads is 72%, 93%, and 94% for the marine, thermal and gut samples, respectively. These numbers set the maximum percentage of reads that can be assigned by the assembly approaches. Accordingly, the most complete classification is achieved for the gut sample, allowing the assignment of 73% of the reads. A significant reduction is observed for the thermal sample (65% of reads assigned), even if the percentage of mapped reads is almost the same. This must be attributed to representation biases in the database: this sample belongs to a much less studied habitat, and therefore close taxa are less represented in the database, complicating the assignment. Finally, the marine sample is the most difficult to annotate by assembly (51% of the reads), because of the lower percentage of mapped reads.

Logically, the percentage of assignment is higher when using the combination of mapping reads to genes and contigs (Ac). Using the contig annotation can overcome unannotated genes, while using gene annotations are not affected by the lack of consensus needed for contig assignment.

Annotation of the raw reads (RR) resulted in 10-20% less classified reads, with the gut metagenome being the best annotated (65%), and marine and thermal having similar percentages (42-45% annotated reads). Kraken2 classification provides less annotations, as much as 25-30% less than the assembly. Again, the gut metagenome is the one having more assignments, benefiting of the increased completeness of the databases in gut-associated taxa. Finally, Metaphlan2 is able to classify very few reads, which is expected because it only annotates marker (clade-specific) genes.

The relative taxonomic composition at the phylum level obtained by each approach can be also seen in Figure 2, discounting the effect of the unclassified reads. Ideally, we should expect the same composition for all methods for the same metagenome, but instead we see that they diverge substantially. One of the most affected phylum is Cyanobacteria, present in thermal and marine samples. Assembly approaches report lower quantities for this taxon than RR and especially Kraken2, which greatly increases its proportion in the two datasets to unrealistic values, particularly in the case of the marine sample. The gene marker approach of MetaPhlan2 predicts less Cyanobacteria than the rest in the marine sample, but much more than the others in the thermal sample. The rest of the taxa are affected in different ways. Rare taxa such as Armatimonadetes in the thermal sample are obtained in greater abundance by the assembly approaches, and ignored by Kraken2 and Methaphlan2, probably because of the absence of complete genomes belonging to this phylum. This is an example of how the gaps in the representation of taxa in the set of available complete genomes can hamper the annotations of methods based on them [9, 27].

While the inferred composition of the gut metagenome is roughly the same for all approaches, the marine and thermal metagenomes vary slightly between raw reads and assembly, and greatly for Kraken2 and Metaphlan2. The thermal metagenome shows big variations that affect for instance the determination of the most abundant taxon in the sample (Chloroflexi by assembly, Proteobacteria by raw reads, and Cyanobacteria by Kraken2 and Metaphlan2). Therefore, the choice of the method can influence greatly the ecological inferences obtained from the analysis.

Next, we compared the discrepancies between the assignments done by different methods, by counting the cases in which the annotations were different at the phylum level (not annotated reads were not considered). The results can be seen in figure 3. Consistently with the previous results, the thermal dataset is the one having more differences. The differences between the assembly methods are rather low, less than 1% of the annotated reads for the thermal metagenome, and almost non-existent for the other samples. Between RR and assembly methods, the differences are also rather low (less than 3% of the reads in the thermal dataset, less than 1% in the others). On the contrary, the differences were much bigger between Kraken2 and the rest: more than 8% for the thermal dataset, more than 4% for the marine, and almost 4% for the gut. This indicates again that the Kraken2 assignment is more dissimilar than the rest.

**Figure 3:**
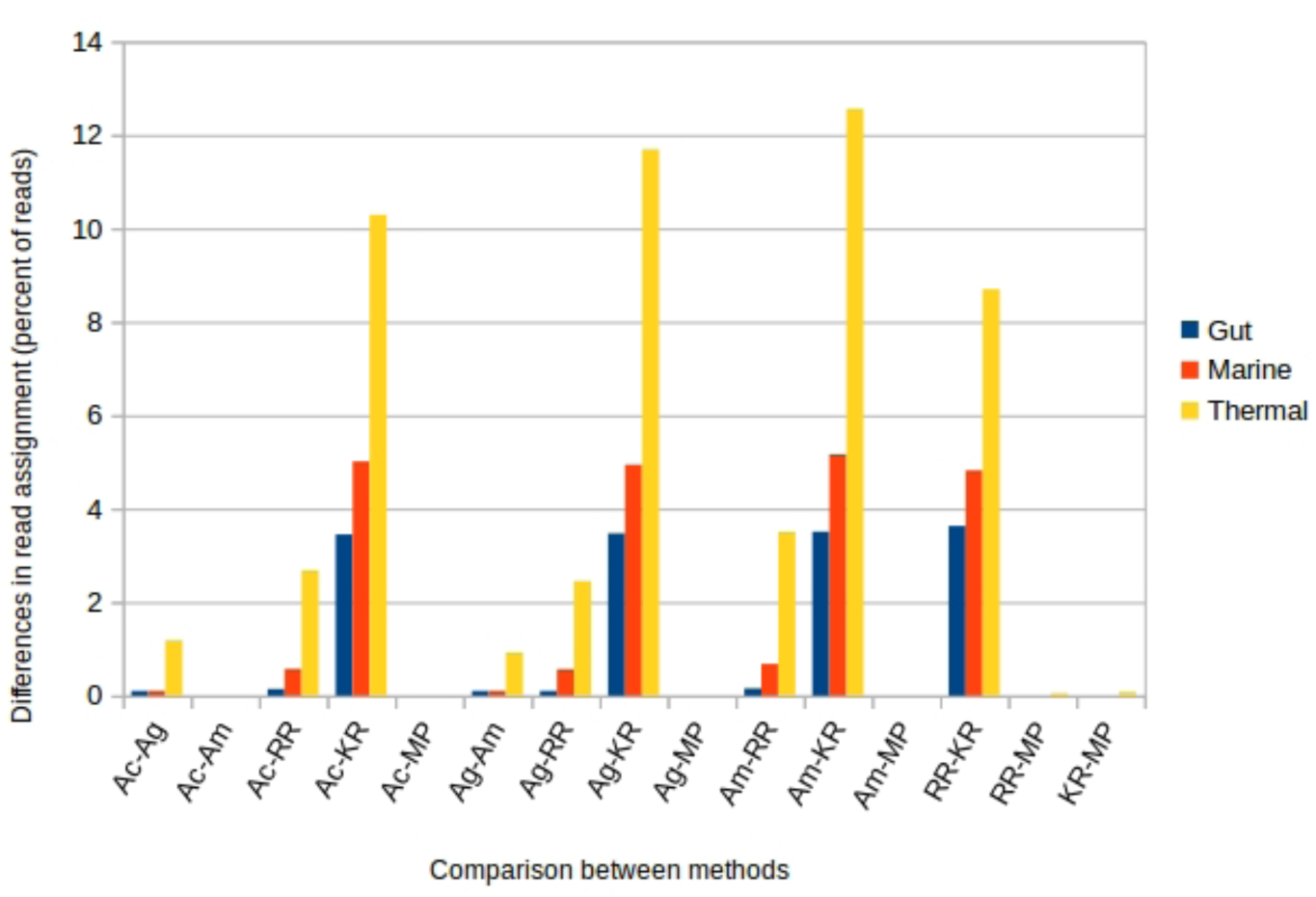
Percentage of discordant assignments between the different methods. Only reads that were classified by both compared methods are considered (i.e. unclassified reads by either method are excluded). A: Assignment by Megahit assembly mapping to: (g: genes; c: contigs; m: combination of contigs and genes). RR: Assignment by raw reads; KR: Kraken2; MP: Metaphlan2.

Then we studied the distribution of functions by the assignment of reads to KEGG functions with the Ag (F_Ag) and RR (F_RR) approaches. Kraken2 and Metaphlan2 were skipped since they do not provide functional annotation, and Ac and Am because there is not contig annotation for functions (each gene has a different function). F_Ag is again able to classify more reads than the F_RR in all metagenomes, even if the difference is small (Figure 4 top). In contrast, F_RR assignment is able to detect a much higher number of KEGGs in all cases (Figure 4 bottom). These correspond to low-abundance functions. The percentage of functions represented by less than 10 reads in F_RR is 20%, 15% and 23% for marine, thermal and gut metagenomes, respectively. These could correspond to low-coverage parts of the metagenome that were not assembled.

**Figure 4:**
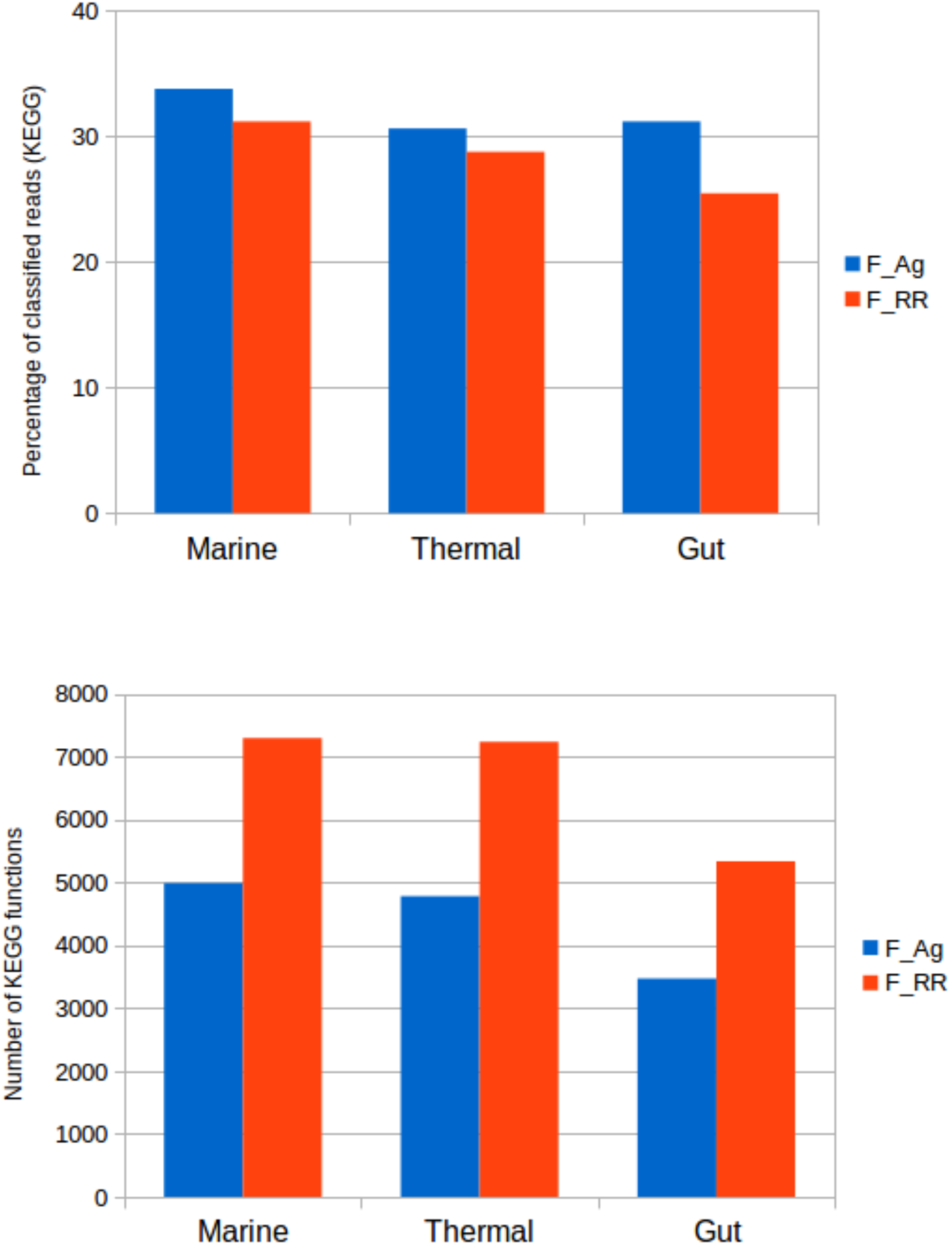
Top, percentage of reads classified in KEGG functions, by raw reads and assembly approaches, in the three metagenomes. Bottom, number of KEGG functions.

Figure 5 (left) shows the distribution of abundances of each KEGG function as rank-abundance curves. Distributions for F_Ag and F_RR are almost indistinguishable, except for the higher number of KEGG IDs detected by raw reads, and the slightly higher abundance for all functions using the assembly, because of the higher number of reads assigned by this method. A comparison of the abundance of KEGG functions can be seen in figure 5 (right), where the good fitting indicates that there are not big differences between the functional assignments by both methods.

**Figure 5:**
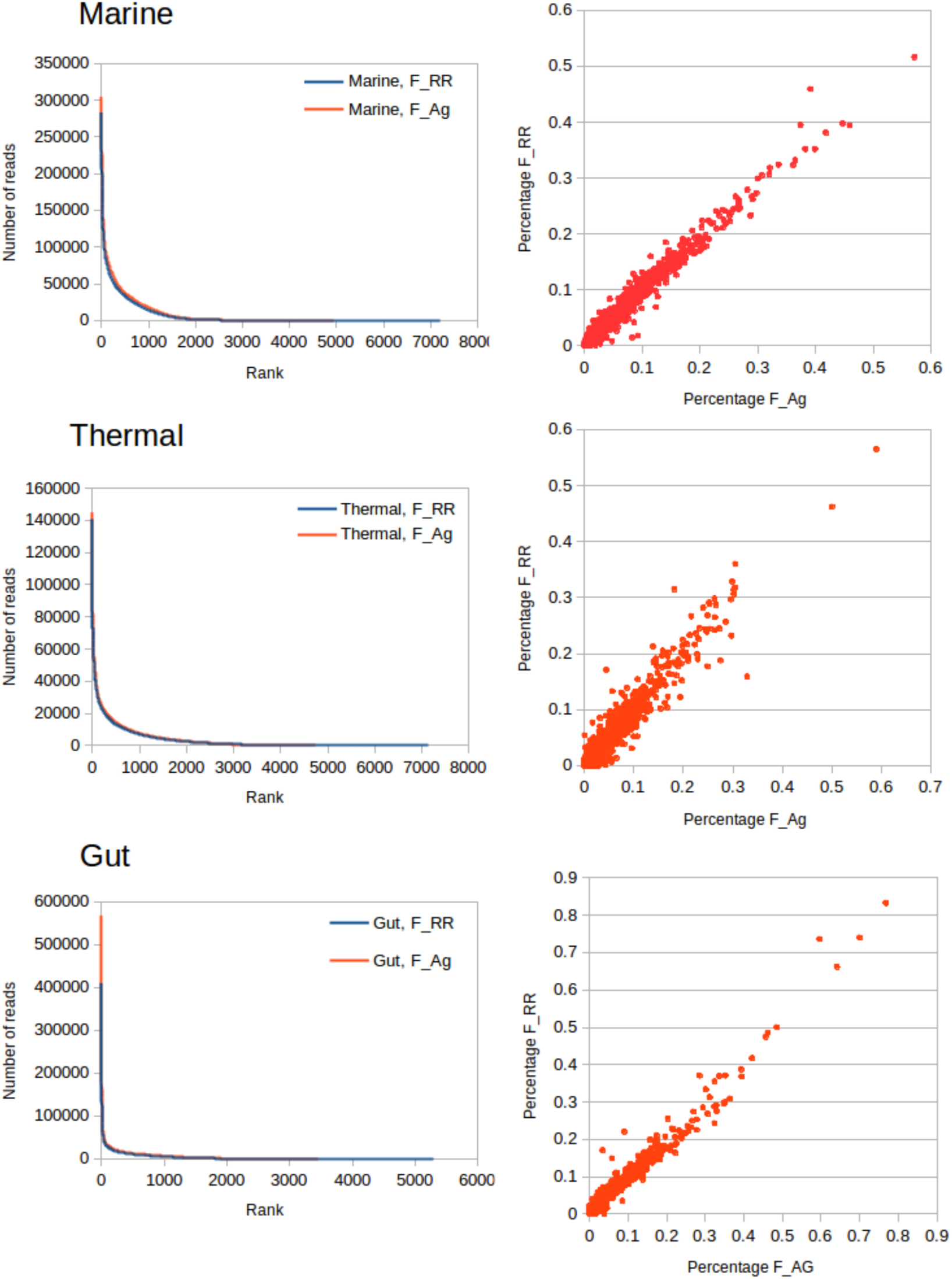
Comparison of functional assignments for all metagenomes. Left: Rank/abundance curves for the distribution of KEGG functions the three metagenomes, classifying either by raw reads (F_RR) or by assembly and mapping to genes (F_Ag). Right: Scatter plots showing the abundance percentages for each KEGG function for both approaches.

### Mock communities

To better estimate the performances of each method of assignments, we created mock communities simulating microbiomes of marine, thermal, and gut environments. We selected 35 complete genomes from species known to be associated to these environments, according to [26], and created mock metagenomes by selecting 1 million (1M) reads from them, in variable proportions. The composition of these mock metagenomes can be found in Supplementary File 1. We analysed these metagenomes using the same methods than for real metagenomes. The results for the phylum rank can be seen in figure 6, and for the family rank in supplementary figure 2.

**Figure 6:**
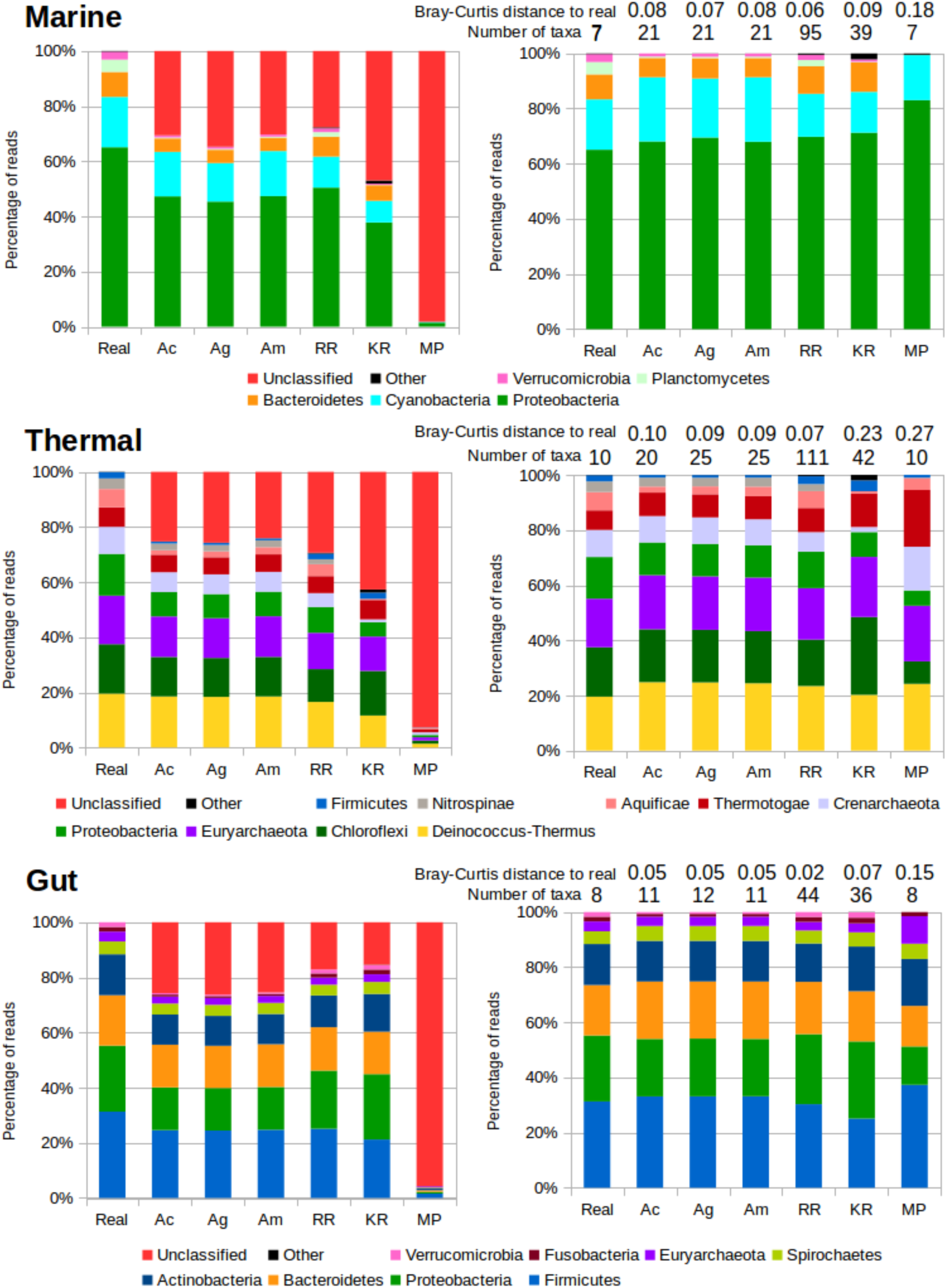
Taxonomic assignments for the mock communities. Left panels show the results for all the reads, right panels show the results removing unclassified reads and scaling to 100%. Real: Real composition of the mock community. Ac, Assembly and mapping reads to contigs. Ag, Same but mapping reads to genes. Am, same but mapping genes first to contigs, then to genes. RR, raw reads assignment. KR: Kraken2. MP: Metaphlan2. Numbers above the bars in the right panels correspond to the Bray-Curtis distance to the composition of the original microbiome, and the number of taxa (phyla) recovered by each method, with the real number of taxa present in the mock metagenome indicated in the “Real” column.

The methods classifying more reads is RR for marine, Am for thermal, and Kraken2 for gut. As expected, the assembly approaches work better when the assemblies are more complete (The percentage of mapped reads in the assemblies is 75%, 84% and 81% for marine, thermal and gut, respectively), while Kraken2 seems to be especially suited to classify gut metagenomes, but misses many reads for metagenomes from other environments. RR also classifies more reads for gut metagenomes, indicating that database completion, which is higher for gut genomes, is an important factor. When removing the unclassified reads, the resulting composition is variable between approaches. We measured the Bray-Curtis distance to the real composition of the mock metagenome to evaluate the closeness of the observed results to the expected ones. The results are rather close to the original composition for the assembly approaches and RR, with best results for the gut metagenome. Kraken2 performs well for the marine and gut metagenomes, even if it misses entire phyla in some instances (for example, Nitrospinae in the thermal metagenome). Metaphlan2 provides the more distant profile in all cases, also missing some phyla entirely.

We also inspected the number of reported phyla by each method. Excess of phyla will be produced by incorrect assignments. Metaphlan2 is the only method that reports the exact number of phyla in all the mock microbiomes. The assembly approaches provide a few phyla more. RR and Kraken2 report a higher number of superfluous taxa. Especially RR produces a very inflated number (more than ten times higher for the thermal mock microbiome). The used version of Kraken2 provided a maximum of 42 phyla for training, therefore this is the maximum number of phyla that can be obtained. In all cases the number is close to this top.

We next measured the error by inspecting the accuracy of the annotation of the individual reads (figure 7). All methods perform well for the gut metagenome, with less that 1% error at the phylum rank, and also at the family rank (except for Kraken2). Nevertheless, substantial differences appear for the other two environments, where errors increase significantly. At phylum rank, the most mistakes are done for the thermal metagenome, while at family rank, the marine metagenome is the most challenging. This is unrelated to the number of taxa in both metagenomes, as the thermal one has more phyla and families. According to that criterion, the most precise method is Metaphlan2, that makes no errors, although we must consider the low number of reads that can be used for classification with this method. Since it also provides a skewed composition, we can conclude that this method performs differently for the diverse taxa. The assembly methods have less that 1% error in all cases, and annotation by contigs is more accurate than by genes, evidencing the advantage of having contextual information of the rest of the genes. RR annotation increases significantly the error rate of the assemblies, reaching 4% for the thermal metagenome at the family level. Kraken2 is the method making more errors, more than 4% for thermal and marine metagenomes at the phylum level, and reaching more than 10% for the marine metagenome at the family level. This is also reflected in the high amount of “Other taxa” classifications for Kraken2 in the figure 6.

**Figure 7:**
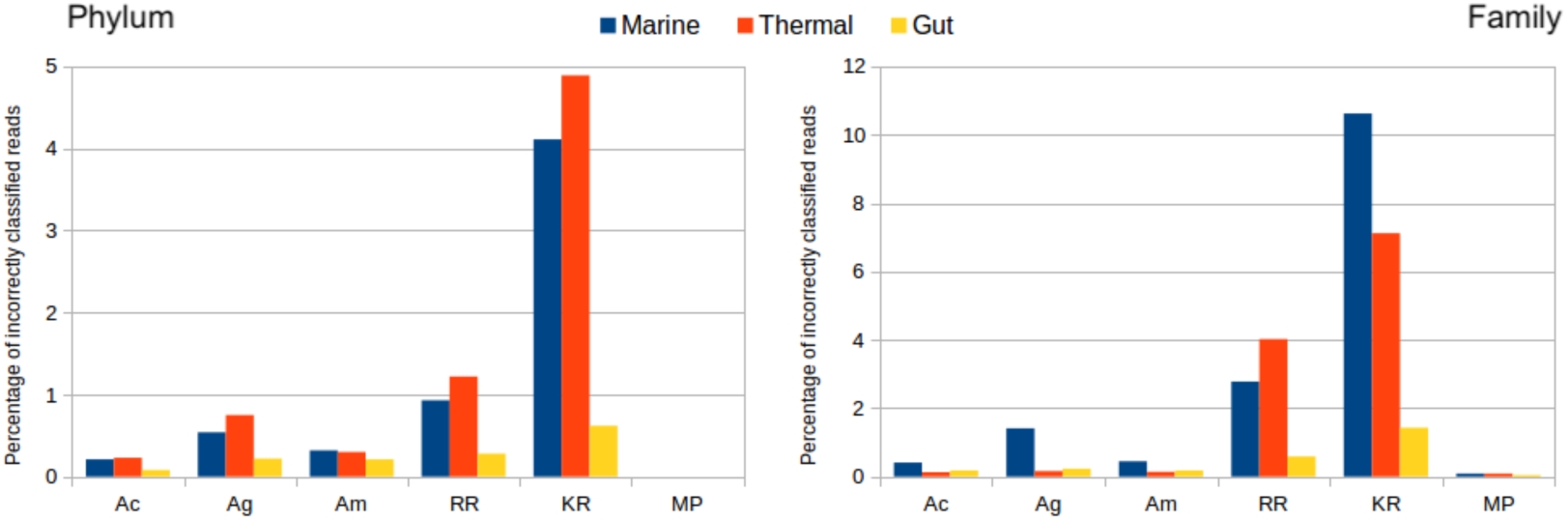
Percentage of incorrectly assigned reads, where the assignment is discordant with the known origin of the read. Left panel, phylum taxonomic rank. Right panel, family taxonomic rank. Ac, Megahit assembly and mapping reads to contigs. Ag, Same but mapping reads to genes. Am, same but mapping genes first to contigs, then to genes. RR, raw reads assignment. KR: Kraken. MP: Metaphlan2.

We were aware that our results could be dependent on metagenomic size, especially those related to the assemblies for which the number of sequences is a critical factor. Therefore, we did additional tests to evaluate the performance of each method regarding the size of the metagenome. Our hypothesis was that methods that classify reads independently (RR, Kraken2 and Metaphlan2) would not be influenced, while the annotation by assembly could be seriously impacted. We created several mock metagenomes of different sizes for marine, thermal and gut environments, extracting reads from genomes strongly associated with these environments [26]. We created mock metagenomes for 2K, 5K, 1M, 2M and 5M paired sequences, all with the same composition of species (Supplementary table 1). We annotated these datasets using the different methods, and calculated the Bray-Curtis distance between the resulting distribution of taxa and the real one. The results can be seen in figure 8 for the phylum rank, and in Supplementary figure 3 for the family rank.

**Figure 8:**
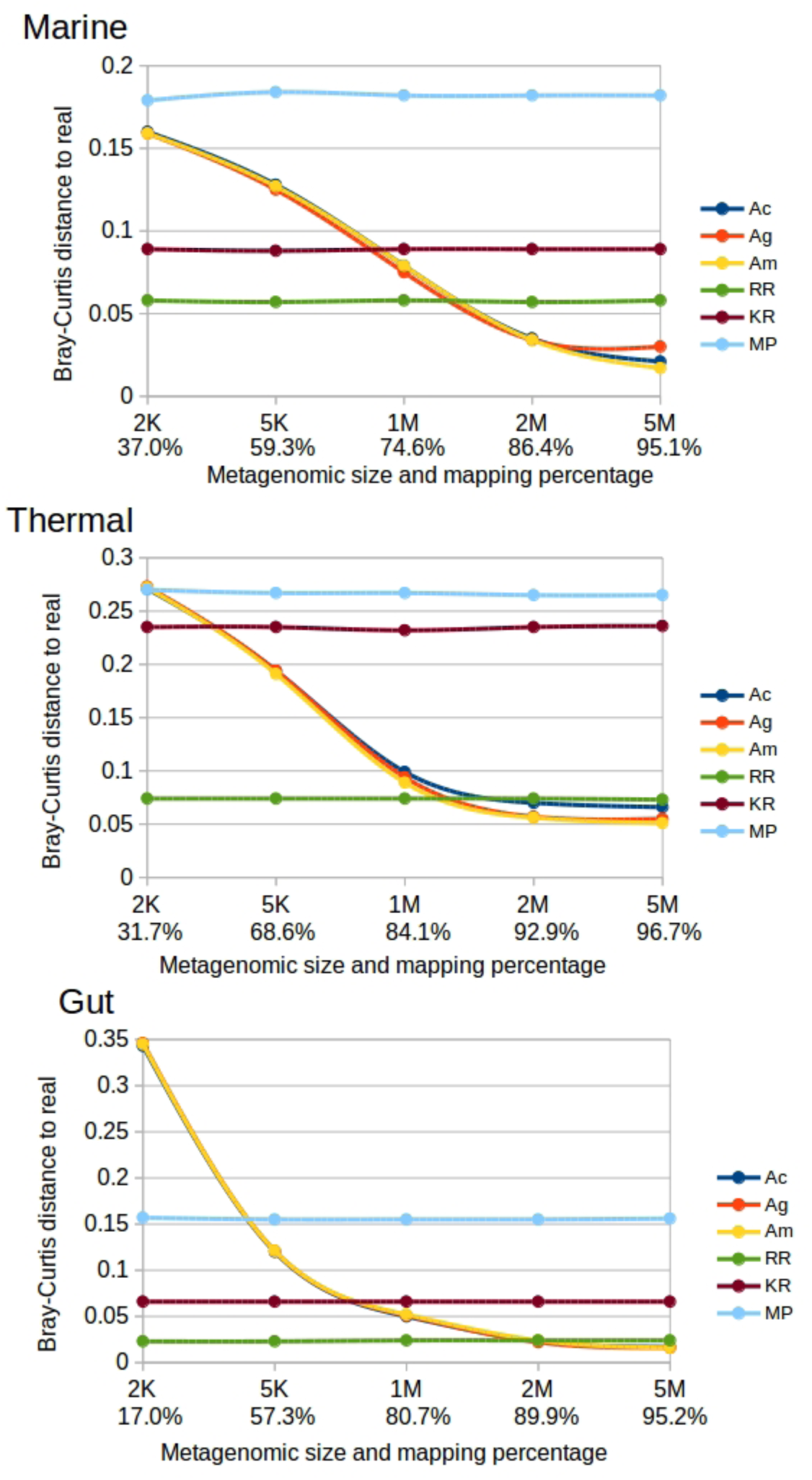
Bray-Curtis distance to the real composition of the marine mock community, for several sample sizes, at phylum rank. Ac, Megahit assembly and mapping reads to contigs. Ag, Same but mapping reads to genes. Am, same but mapping genes first to contigs, then to genes. RR, raw reads assignment. KR: Kraken2. MP: Metaphlan2.

As we expected, RR, Kraken2 and Metaphlan2 are not affected by the size of the metagenomic sample. Metaphlan2 is the method diverging more from the actual composition, except for the thermal mock community at family rank, where Kraken2 is the farthest. Of these three methods directly assigning reads, RR is clearly the most accurate. Again, these methods perform much better for the gut mock metagenome than for the rest.

The assembly methods are highly dependent of the amount of reads that can be assembled. For very small samples, where less than 50% of the reads in the assembly, it provides much more divergent classifications than other methods. When the percentage of assembled reads is in the range of 80-90%, they obtain similar results than RR. When the percentage of assembled reads is higher than that, annotation by assembly outperforms the other methods. This indicates that the coverage of the metagenome, which is directly related to the percentage of assembled reads, can be seen as the factor determining if it is more advantageous using RR or assembly methods for analysing metagenomes.

We also analysed the functional assignment of these mock metagenomes. We used the assignment by homology of full genes to KEGG functions as a reference, and classified the reads using the Assembly (F_Ag) and Raw Read (F_RR) annotation, as above. The results can be seen in the figure 9

**Figure 9:**
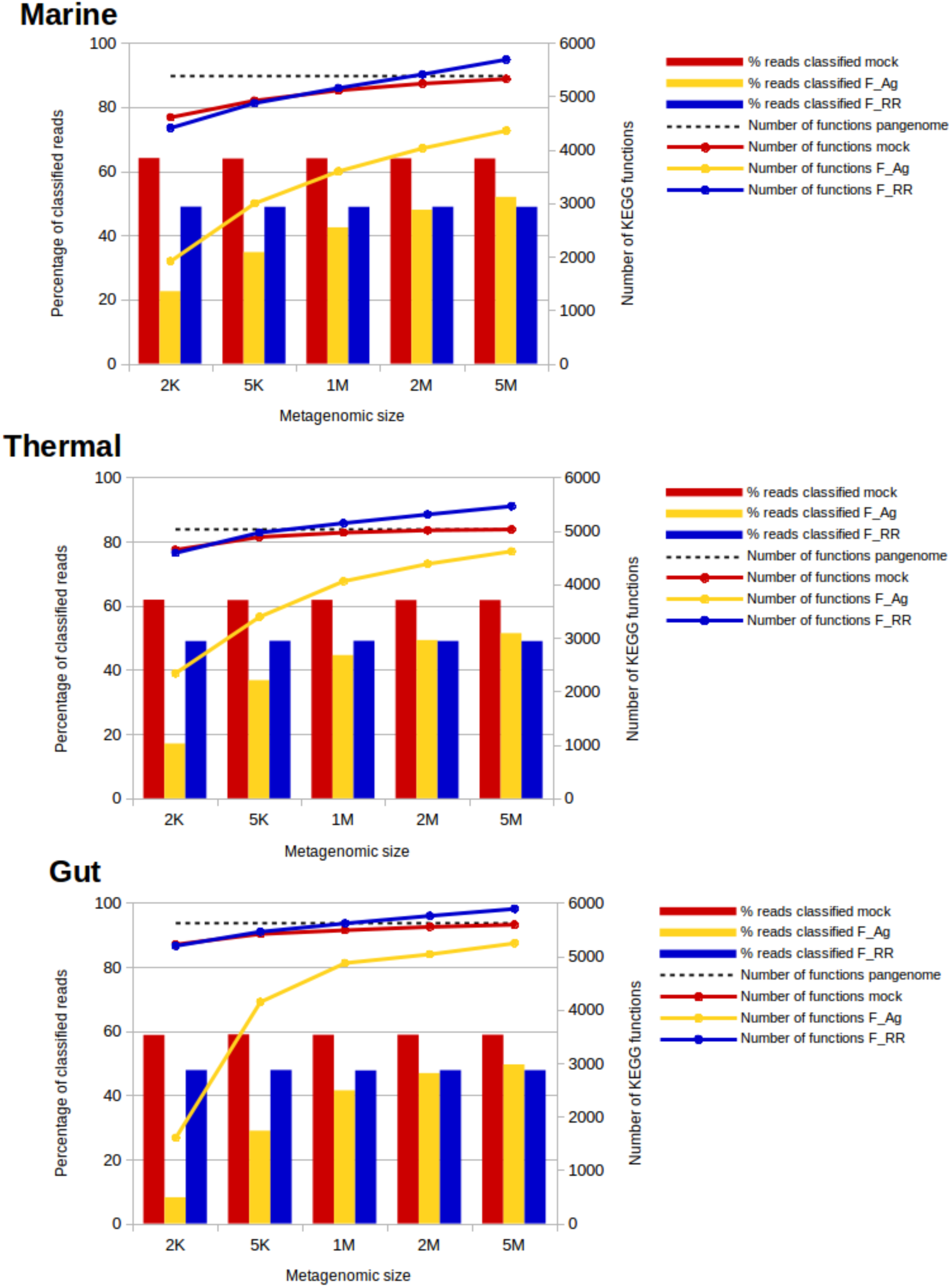
Percentage of reads classified to KEGG functions (bars) and number of KEGG functions (lines) provided by the different functional assignment methods, for the different sizes of the mock metagenomes. The data labelled as “mock” correspond to the annotations of the reads based on the original annotations of the genomes. The data corresponding to “pangenome” correspond to the total number of functions in all the genomes used to build the mock communities.

The percentage of reads that can be classified is around 60% for all metagenomes. The rest correspond to reads from genes with no known function or with no KEGG associated. RR classification is around 50% in all cases. It does not vary with metagenomic size because for each size the reads are extracted from the same background distribution of functions and they are annotated independently. F_Ag functional assignment, in turn, varies with size since it depends on the completion of the assembly, as stated above. We can see that for the biggest sizes, the percentage of assignments is larger for F_Ag than for F_RR. Therefore, F_RR is advantageous for small metagenomes, while assembly excels at bigger sizes. In this case there is no evident differences regarding the diverse environments.

Concerning the number of functions detected, we can see how the number is variable, reflecting the fact that rare functions are easily detected when metagenomes are bigger, but this number tends to stabilize, ultimately reflecting the total number of functions in the pangenome. It can be seen how the F_RR approach is over-predicting the number of functions, exceeding these actually present in both the metagenome and the pangenome, thus producing false positives. Even worse, the number of predicted functions increases linearly and shows no saturation, in contrast to the real number of functions. On the other hand, F_Ag produces a very low number of functions when the metagenomes are small, but it quickly increases to numbers close to the real ones for bigger sizes.

We also quantified the number of wrong annotations by comparing the annotation of reads by each method with regard to the real one. The results can be seen in figure 10.

**Figure 10.**
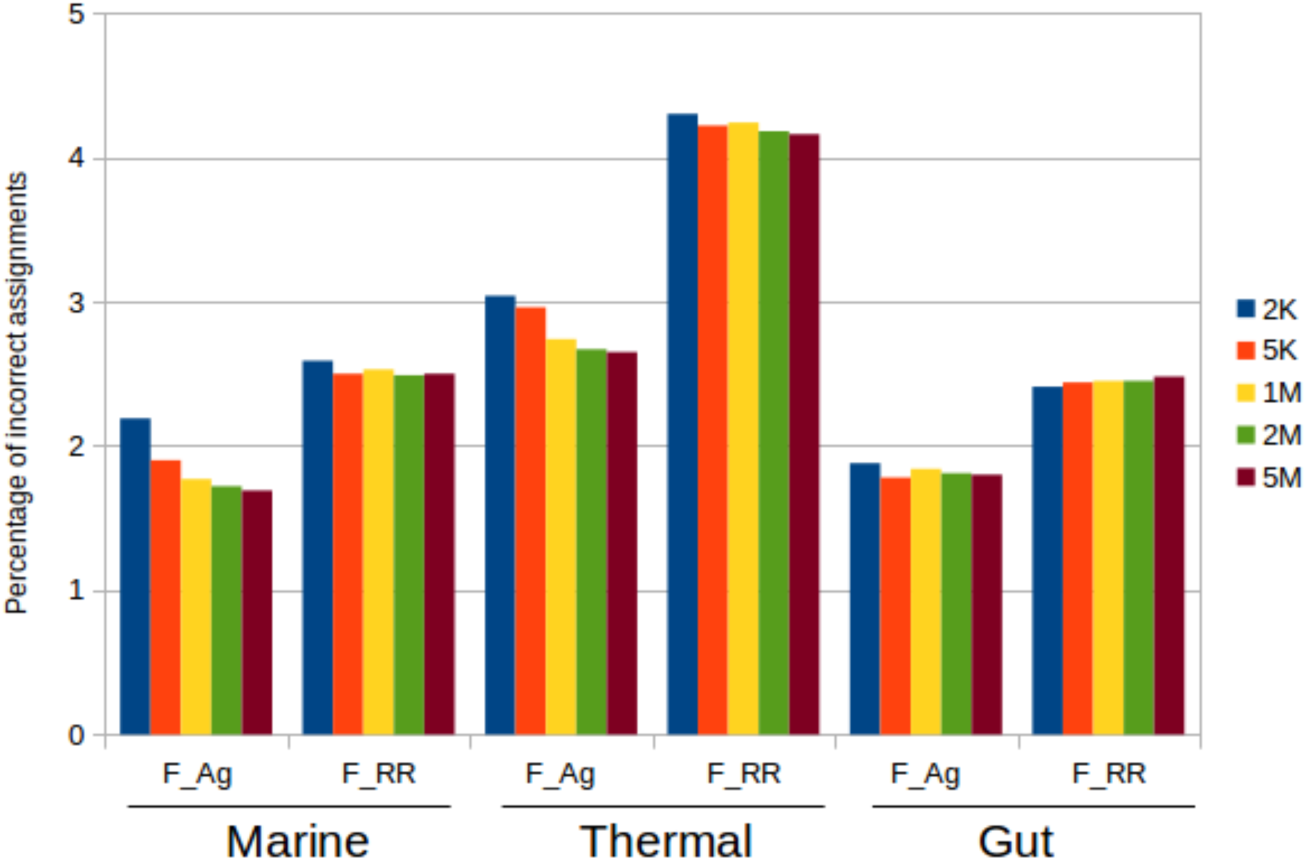
Percentage of incorrect assignments for F_Ag and F_RR in each of the mock metagenomes. A read is incorrectly annotated when the provided function is different than the one from the original gene in the genome.

F_RR assignments are always more error-prone. As for the taxonomic analysis, the thermal metagenome is the most difficult to annotate, and the gut one the easiest. The percentage of errors does not vary with sizes, and it is above 4% in the thermal metagenome. Apparently, metagenomes from this environment are more difficult to classify, in accordance with the previous results for true metagenomes. The F_Ag annotations are more precise, not exceeding the threshold of 3% errors. The influence of sizes can be noticed also here, with usually fewer errors in the bigger metagenomic sizes, but it is not so evident. For instance, the gut example shows a very stable error rate around 1.8%, irrespectively of the metagenomic size.

## Discussion

Different approaches can be used for the taxonomic and functional annotation of metagenomes. Working with raw reads, its annotation can be done using homology, k-mer composition or gene marker searches. But we also can assemble the reads and use the assembly as a framework for the annotation, since this will provide longer fragments of genes (or complete ones) and contextual information. There is not a standard way of proceed, and examples of each approach can be found in the literature. However, it is unclear how the diverse approaches can influence the accuracy and reproducibility of the results. We wanted to explore the characteristics of each method to, if possible, provide hints helping the choice of the most appropriate method.

Assembly and especially the assignment of raw reads are demanding methods in time and computational resources. In contrast, Kraken2 and Metaphlan2 are very fast methods that can be very useful to obtain a quick view of the diversity of the metagenomes. Nevertheless, the analysis of mock metagenomes shows that these methods are less accurate, especially for non-human-associated environments. They are rather sensitive to the composition of the databases, and their performance decreases when rare or less species are present in the metagenome. This latter drawback also holds true for the assignment of raw reads by homology.

While for the methods above database completeness is the main factor determining their performance, for the assembly approaches the critical issue is sequencing depth, that in turn influences the completeness of the assembly. When the assemblies are complete enough to recruit 85% or more of the original reads, assembly approaches are more advantageous in terms to percentage of reads classified, smaller number of errors and importantly, similarity to the real scenario. When sequencing depth is reduced because of both a high microbial diversity and a small number of reads, the assignment of raw reads could be preferred. Assembly approaches seem to be less influenced than others by database completeness because having longer sequences (full genes instead of short reads) is advantageous when only distant homologies can be found, and, for taxonomic annotation, having the contextual information of the contigs helps to infer annotations for all genes on it. Nevertheless, they are also affected to some extent by database composition.

Therefore, when dealing with small metagenomes from well-studied ecosystems, such as these human-associated, the usage of raw read assignment can be advantageous for taxonomic assignments. Otherwise assembly approaches should be preferred. This is especially true if we want to obtain bins, where co-assembly of metagenomes is mandatory. We did not consider the effect of co-assemblying in the annotation, but since it helps to obtain more and longer contigs and therefore to map more reads to the assembly, it is expected to improve the annotations even more. It would be also possible to follow a combined approach in which assembly is done and used as a reference, and then the remaining unmapped reads are classified independently.

As for functions, the KEGG assignments for the real metagenomes show a high degree of correlation between assembly and raw read annotation. It is generally harder to annotate the functions than the taxonomic origin, because short reads are often not discriminative enough to distinguish between functions. Read mapping to promiscuous domains that can participate in different proteins or to conserved regions between genes difficults its accurate assignment. Consequently, assembly annotation provides a higher percentage of functional classification. On the other hand, functions represented by a few reads will be probably missed by the assembly approaches. Because of this, raw read assignment provide a higher number of functions than the assembly. Given these advantages and disadvantages of each method, if one is interested in looking for particular, specific functions, a combination of both approaches is advisable.

### Availability of data and materials

The datasets supporting the conclusions of this article are available in the repository http://botero.cnb.csic.es/lab35/suppl/method_comparison

## Acknowledgements

This research was funded by projects CTM2013-48292-C3-2-R and CTM2016-80095-C2-1-R, Spanish Ministry of Economy and Competitiveness.

---

**Suppl Fig 1:**
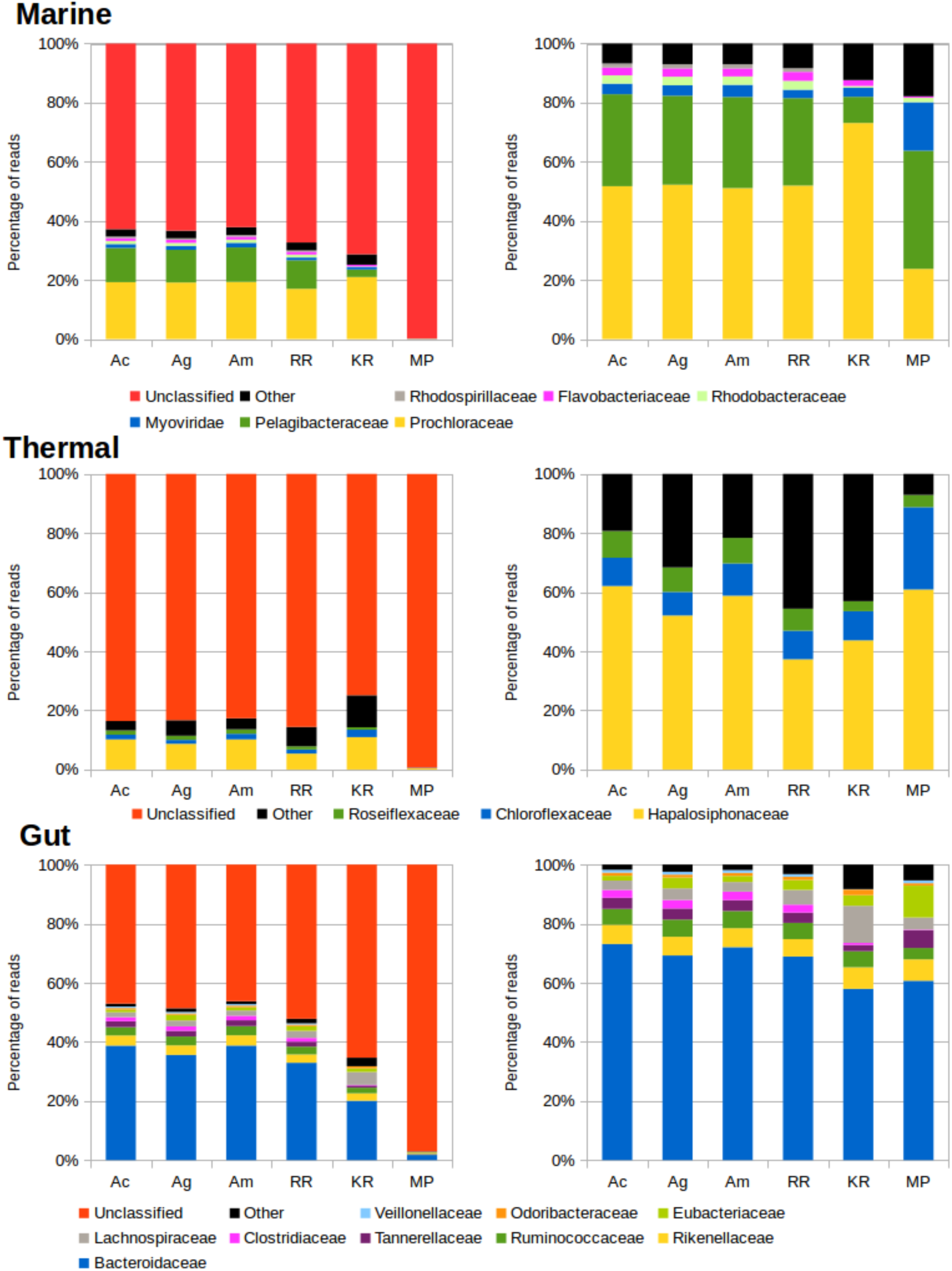
Comparison of read assignments by different methods, at the family rank: Ac, Megahit assembly and mapping reads to contigs. Ag, Same but mapping reads to genes. Am, same but mapping genes first to contigs, then to genes. RR, raw reads assignment. KR: Kraken. MP: Metaphlan2. Left: All reads considered. Right: Discounting unclassified reads.

**Suppl Fig 2:**
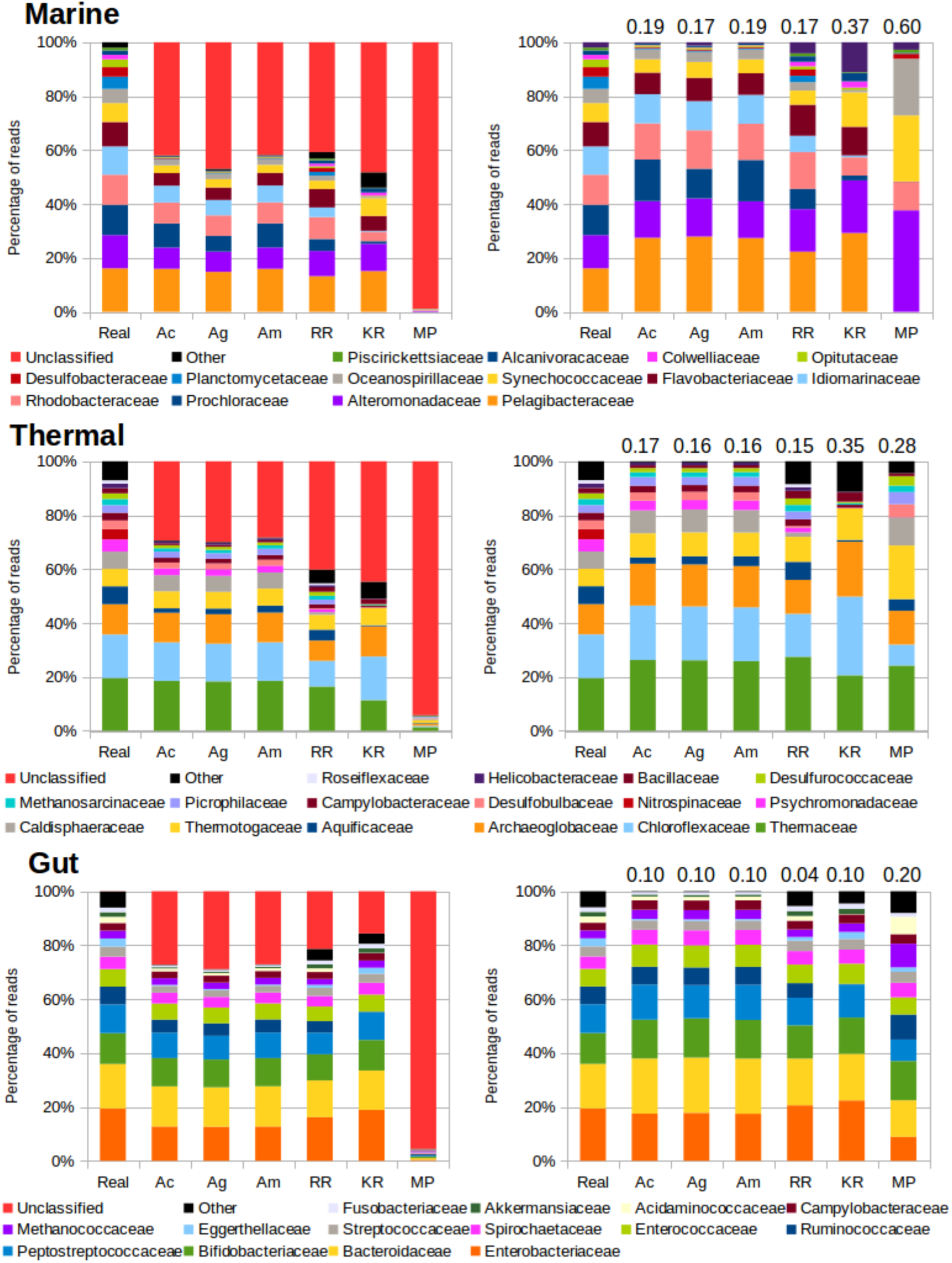
Taxonomic assignments for the mock communities. Left panels show the results for all the reads, right panel, the results removing unclassified reads and scaling to 100%. Real: Real composition of the mock community. Ac, Megahit assembly and mapping reads to contigs. Ag, Same but mapping reads to genes. Am, same but mapping genes first to contigs, then to genes. RR, raw reads assignment. KR: Kraken. MP: Metaphlan2. Numbers above the bars in the right panels correspond to the Bray-Curtis distance to the composition of the original microbiome.

**Suppl figure 3:**
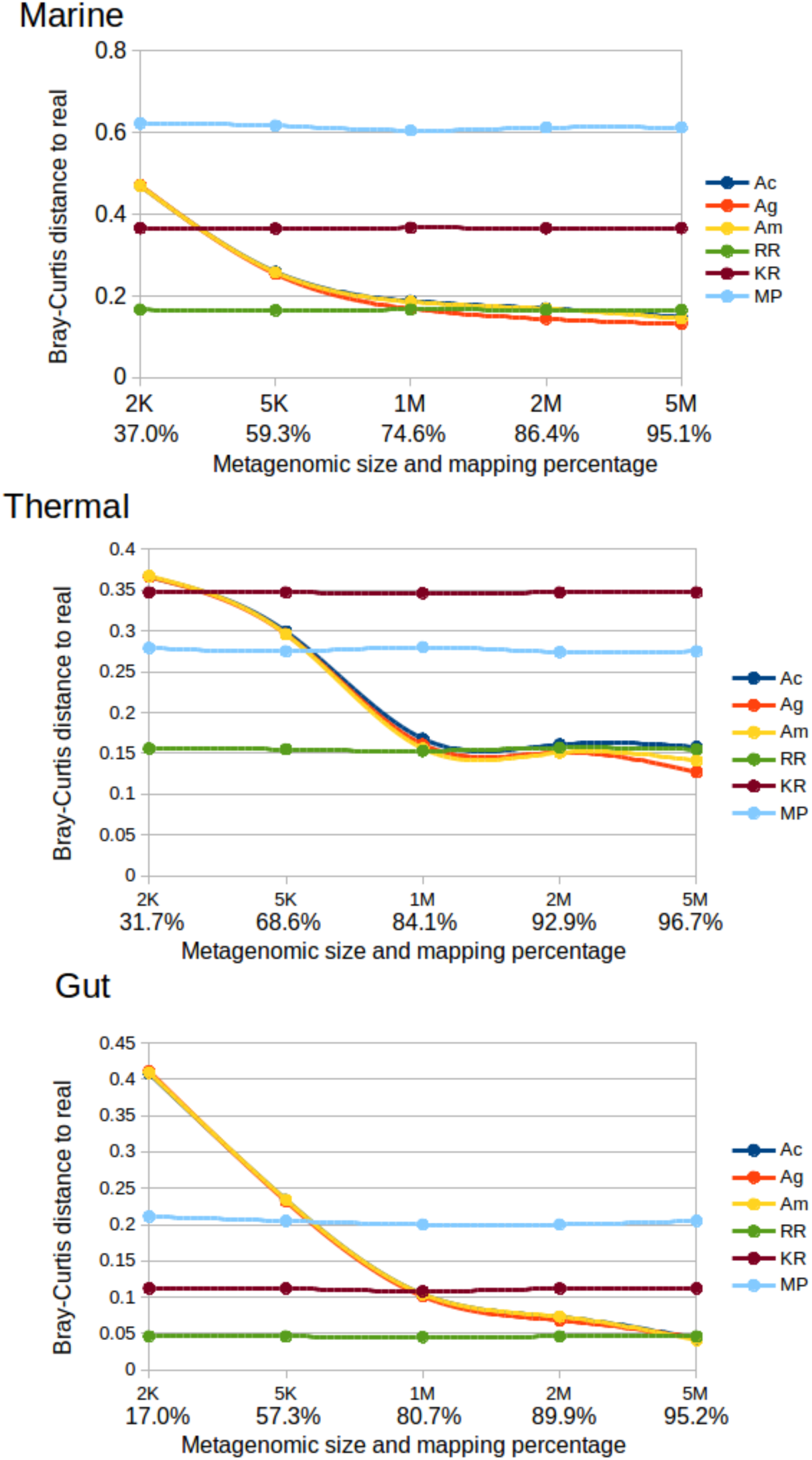
Bray-Curtis distance to the real composition of the marine mock community, for several sample sizes, at family rank. Ac, Megahit assembly and mapping reads to contigs. Ag, Same but mapping reads to genes. Am, same but mapping genes first to contigs, then to genes. RR, raw reads assignment. KR: Kraken2. MP: Metaphlan2.

